# Blind tests of RNA nearest neighbor energy prediction

**DOI:** 10.1101/052621

**Authors:** Fang-Chieh Chou, Wipapat Kladwang, Kalli Kappel, Rhiju Das

**Author notes:** Correspondence to: Rhiju Das.

## Abstract

The predictive modeling and design of biologically active RNA molecules requires understanding the energetic balance amongst their basic components. Rapid developments in computer simulation promise increasingly accurate recovery of RNA’s nearest neighbor (NN) free energy parameters, but these methods have not been tested in predictive trials or on non-standard nucleotides. Here, we present the first such tests through a RECCES-Rosetta (<u>R</u>eweighting of <u>E</u>nergy-function <u>C</u>ollection with <u>C</u>onformational <u>E</u>nsemble <u>S</u>ampling in Rosetta) framework that rigorously models conformational entropy, predicts previously unmeasured NN parameters, and estimates these values’ systematic uncertainties. RECCES-Rosetta recovers the ten NN parameters for Watson-Crick stacked base pairs and thirty-two single-nucleotide dangling-end parameters with unprecedented accuracies – root-mean-square deviations (RMSD) of 0.28 kcal/mol and 0.41 kcal/mol, respectively. For set-aside test sets, RECCES-Rosetta gives an RMSD of 0.32 kcal/mol on eight stacked pairs involving G-U wobble pairs and an RMSD of 0.99 kcal/mol on seven stacked pairs involving non-standard isocytidine-isoguanosine pairs. To more rigorously assess RECCES-Rosetta, we carried out four blind predictions for stacked pairs involving 2,6-diaminopurine-U pairs, which achieved 0.64 kcal/mol RMSD accuracy when tested by subsequent experiments. Overall, these results establish that computational methods can now blindly predict energetics of basic RNA motifs, including chemically modified variants, with consistently better than 1 kcal/mol accuracy. Systematic tests indicate that resolving the remaining discrepancies will require energy function improvements beyond simply reweighting component terms, and we propose further blind trials to test such efforts.

**Significance:** Understanding RNA machines and how their behavior can be modulated by chemical modification is increasingly recognized as an important biological and bioengineering problem, with continuing discoveries of riboswitches, mRNA regulons, CRISPR-guided editing complexes, and RNA enzymes. Computational strategies to understanding RNA energetics are being proposed, but have not yet faced rigorous tests. We describe a modeling strategy called ‘RECCES-Rosetta’ that models the full ensemble of motions of RNA in single-stranded form and in helices, including non-standard nucleotides such as 2,6-diaminopurine, a variant of adenosine. When compared to experiments, including blind tests, the energetic accuracies of RECCES-Rosetta calculations are at levels close to experimental error, suggesting that computation can now be used to predict and design basic RNA energetics.

## Introduction

RNA plays central roles in biological processes including translation, splicing, regulation of genetic expression, and catalysis (1, 2) and in bioengineering efforts to control these processes (3–5). These critical RNA functions are defined at their most fundamental level by the energetics of how RNA folds and interacts with other RNAs and molecular partners, and how these processes change upon naturally occurring or artificially introduced chemical modifications. Experimentally, the folding free energies of RNA motifs can be precisely measured by optical melting experiments, and a compendium of these measurements have established the nearest neighbor (NN) model for the most basic RNA elements, including double helices with the four canonical ribonucleotides (6). In the NN model, the stability of a base pair is assumed to only be affected by its adjacent base pairs, and the folding free energy of a canonical RNA helix can be estimated based on NN parameters for each stacked pair, an initialization term for the entropic cost of creating the first base pair, and corrections for different terminal base pairs. While next-nearest neighbor effects and tertiary contacts are not treated in the NN model (7–9), the current NN model gives accurate predictions for the folding free energies of canonical RNA helices (< 0.5 kcal/mol for helices with 6-8 base pairs) (10, 11) and can be extended to single-nucleotide dangling-ends, chemically modified nucleotides, and more complex motifs, such as non-canonical base pairs, hairpins, and internal loops (11–14). However, it is currently not feasible to experimentally characterize the energetics of all RNA motifs due to the large number of possible motif sequences and the requirement of specialized experiments to address complex motif topologies such as three-way junctions (15–17). These considerations, and the desire to test physical models of RNA folding, have motivated several groups to pursue automated computational methods to calculate the folding free energies of RNA motifs.

Current computational approaches are beginning to recover NN parameters for the simplest RNA motifs with accuracies within a few-fold of the errors of experimental approaches. For example, the Rosetta package has been developed and extensively tested for structure prediction and design of macromolecules, including RNA. Recent successes at near-atomic resolution have leveraged an all-atom ‘score function’ that includes physics-based terms (for hydrogen bonding, van der Waals packing, and orientation-dependent implicit solvation) and knowledge-based terms (for, e.g., RNA torsional preferences) (18). When interpreting the total score as an effective energy for a conformation, simple Rosetta calculations recover the NN parameters for all canonical stacked pairs with an RMSD of less than 0.5 kcal/mol upon fitting two phenomenological parameters, the Rosetta energy scale and a constant offset parameterizing the conformational entropy loss upon folding each base pair ((18); and see below). In parallel, molecular dynamics studies have demonstrated calculation of folding free energies of short RNA hairpins using umbrella sampling, molecular mechanics-Poisson Boltzmann surface area (MM-PB/SA), free energy perturbation, and other methods (19–22). While these calculations have not yet accurately recovered folding free energies (errors > 10 kcal/mol) (21, 22), relative differences of NN parameters between different sequences and other aspects of RNA motif energetics have been recovered with accuracies between 0.6 to 1.8 kcal/mol (22–24). These error ranges are similar or lower than uncertainties of empirically defined NN parameters for most motifs; original NN energy estimates for G-U dangling ends, single nucleotide bulges, and tetraloop free energies have been corrected by > 1 kcal/mol when revisited in detailed studies (25–28). Overall, computational approaches may be ready for calculations of new energetic parameters, including parameters for these uncertain motifs as well as for motifs involving non-standard nucleotides that are being found throughout natural coding and non-coding RNAs (29, 30) or used to engineer new RNA systems (31, 32). However, the predictive power of these methods has not been evaluated through tests on previously unmeasured nearest neighbor parameters. Predictive tests are particularly important as models are increasing in complexity and risk over-training on previously available data.

Here, we report the first blind tests of a method to computationally predict nearest neighbor energetic parameters. The newly measured parameters involve RNA stacked pairs with the non-natural nucleotide 2,6-diaminopurine (D) paired to uracil (Figure 1). To ensure a rigorous comparison, calculations were carried out by one of us (FCC) and subsequently tested in independent experiments by another author (WK). In preparation for this blind test, we developed a <u>R</u>eweighting of <u>E</u>nergy-function <u>C</u>ollection with <u>C</u>onformational <u>E</u>nsemble <u>S</u>ampling in Rosetta (RECCES-Rosetta) framework to calculate free energies based on density-of-states estimation and expected errors from statistical precision, inaccuracies in the nearest neighbor assumption, and uncertainties in the weights of the underlying energy function. Furthermore, to address previous *ad hoc* assumptions used to fit conformational entropy from data, RECCES calculates the conformational entropy of helix and single-stranded states without fitting of additional parameters. These systematic improvements – and calibration based on previously measured NN parameters – ensured that our blind tests carried sufficient power to rigorously establish the accuracy and limitations of nearest neighbor energy calculations that seek to make non-trivial predictions.

**Figure 1.**
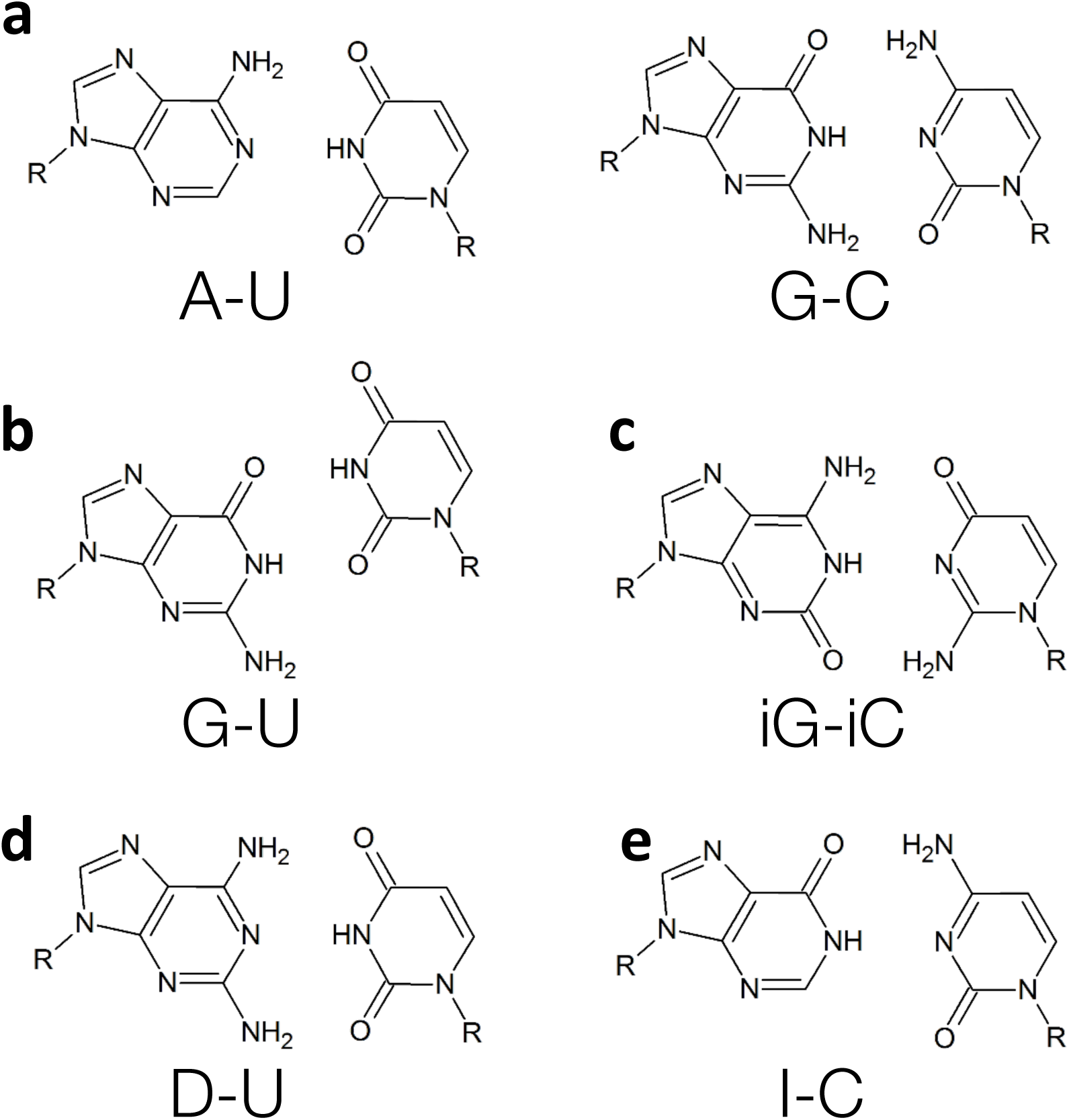
Base pairs involved in nearest neighbor parameters considered in this study. (a) Canonical pairs adenosine-uracil and guanosine-cytidine, (b) guanosine-uracil wobble pair, (c) non-natural isoguanosine-isocytidine, (d) non-natural 2,6-diaminopurine-uracil, and (e) inosine-cytidine.

## Results

In this study, we performed tests of our ability to predict RNA nearest neighbor parameters of the simplest RNA motifs, including previously unmeasured values for chemically modified nucleotides. We first determined whether blind tests would have the power to falsify our energy function by developing a RECCES framework to calculate the expected statistical and systematic errors of our calculation methods and to assess the accuracy of these calculations on previously measured NN parameters. We then carried out blind tests on previously unmeasured NN parameters.

### Recovery of canonical helix and dangling end parameters

Blind tests of a prediction method are not worthwhile if the expected prediction errors significantly exceed the range of possible experimental values – on the order of several kcal/mol for NN parameters. We therefore first sought to determine whether folding free energy calculations with the Rosetta all-atom energy function, previously developed for RNA structure prediction and design, could recover NN energetics for canonical Watson/Crick stacked pairs and whether these calculations’ uncertainties were acceptable for making blind predictions. The Rosetta energy function involves separate component terms for hydrogen bonding, electrostatics, van der Waals interactions, nucleobase stacking, torsional potentials, and an orientation-dependent solvation model. Prior structure prediction and design studies did not strongly constrain the weights of these components (18). Thus, we anticipated that NN parameter prediction would require optimization of the weights and care in uncertainty estimation. To estimate the errors due to weight uncertainties, we sought not a single weight set but instead a large collection of such weights consistent with available data. We hypothesized that making predictions from these different weight sets would let us assess uncertainties in predicting new parameters, enabling us to decide whether it was worthwhile to carry out blind experimental tests.

To discover diverse weight sets consistent with prior NN data, we developed the RECCES framework for sampling conformational ensembles of the single-stranded and helix conformations relevant to NN energy estimation (Figure 2). Through the use of a density-of-states formalism, simulated tempering, and WHAM integration, RECCES allowed the estimation of free energies with bootstrapped errors of less than 0.003 kcal/mol, significantly less than systematic errors of 0.3 kcal/mol (estimated below; see SI Appendix, Tables S1 & S2), using two CPU-hours of computation per molecule. These methods are similar to replica exchange methods in common use in molecular dynamics studies, but are simpler in that they do not require running multiple parallel processes (SI Appendix, Supporting Methods). Importantly, the overall RECCES framework did not require separate fitting of conformational entropy factors, reducing the likelihood of overfitting. Furthermore, starting from these initial simulations, RECCES enabled evaluation of alternative weight sets with negligible additional computation (<0.1 sec) through a rapid reweighting of cached energies. While noisy at low energies (compare green to blue curves in Figure 2c), we confirmed that this reweighting procedure nevertheless led to an acceptable mean calculation error of 0.28 kcal/mol (SI Appendix, Table S3), significantly smaller than the several kcal/mol range of experimental NN parameters (Table 2). Further tests of the nearest neighbor assumption, based on simulations with different helix contexts for each stacked pair, also gave systematic errors of 0.2-0.3 kcal/mol (SI Appendix, Table S1). Hereafter, we conservatively describe the systematic errors of the RECCES-Rosetta NN parameter estimates to be the higher value in this range, 0.3 kcal/mol.

**Figure 2.**
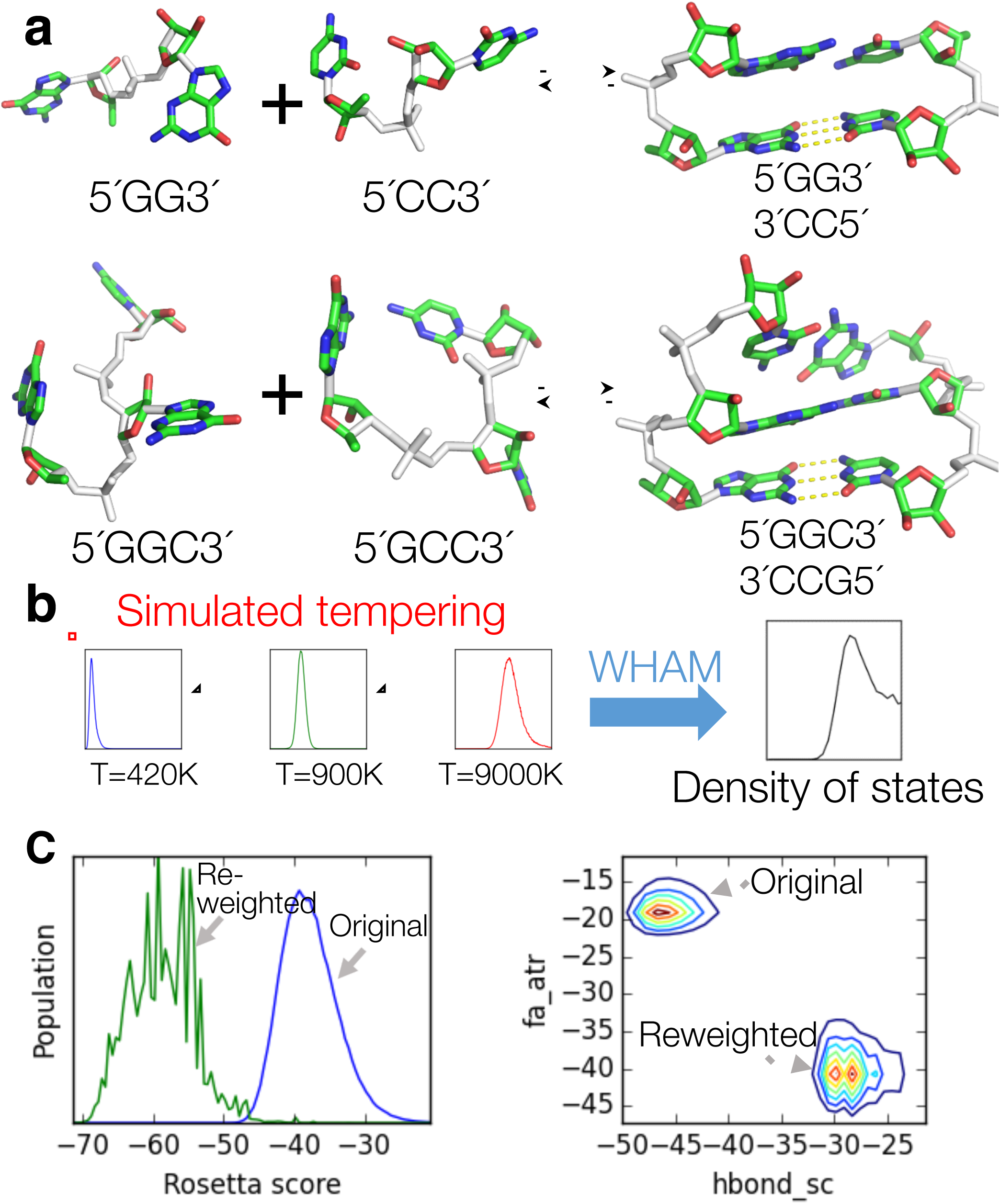
RECCES thermodynamic framework and reweighting. (a) Example systems simulated for this study. Degrees of freedom sampled are colored in white. The relative orientation of first base pair in each helix was fixed (yellow dashes in right-hand panels). The upper and lower panels show the folding reaction two-base-pair and three-base-pair systems, respectively. (b) Density of state estimation by simulated tempering and WHAM. (c) Reweighting demonstration. Left: state population at room temperature before (blue) and after (green) reweighting. Right: 2D population histograms of *fa_atr* (Lennard-Jones attraction) vs. *hbond_sc* (hydrogen bonds) energy components, before and after reweighting.

**Table 2.**
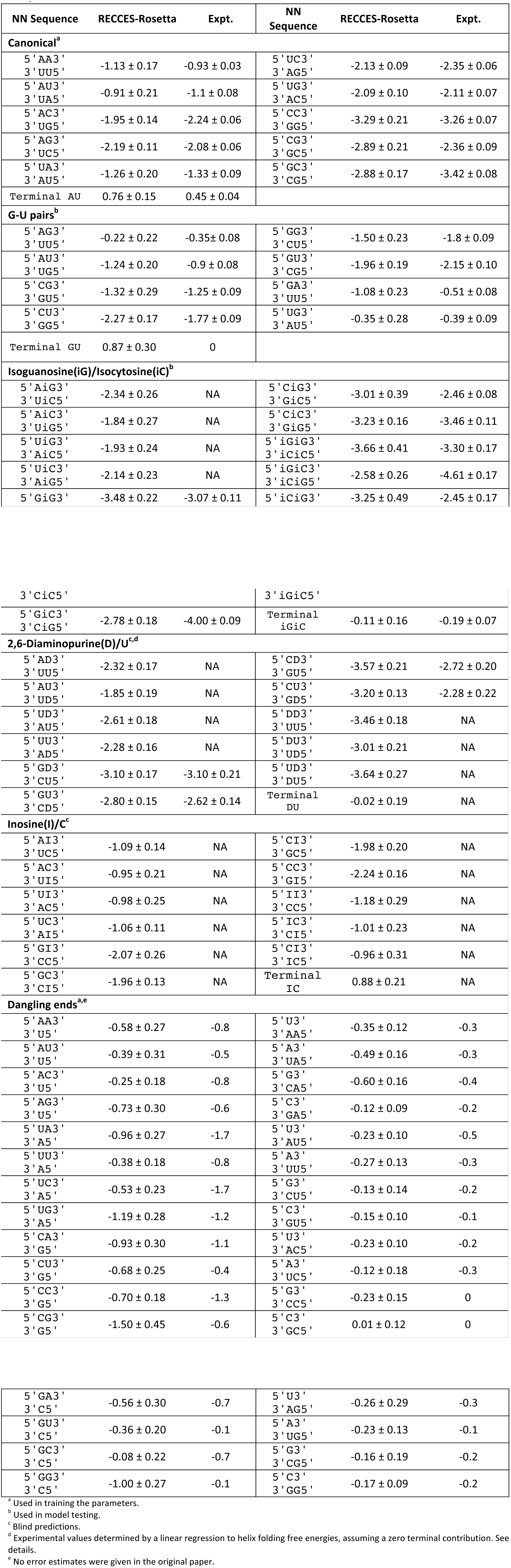
**Experimental NN parameters(27, 36, 49)and RECCES-Rosetta predictions. All values in kcal/mol**.

To obtain a collection of weight sets, we used RECCES to optimize the weights of all terms in the Rosetta score function over numerous runs with different initial values. These optimization runs minimized the mean square error with respect to the NN parameters of 10 canonical stacked pairs (4 base pairs next to 4 base pairs, removing symmetric cases), 32 single-nucleotide dangling-ends (4 nucleotides at either the 5´ or 3´ end of 4 base pairs), and the terminal penalty for A-U vs. G-C. The resulting 9,544 minimized weight sets were highly diverse, even after discarding the weight sets with 5% worst RMSD agreement to training data (SI Appendix, Table S4, describes score terms and summarizes mean and standard deviations of weights; SI Appendix, Table S5, gives five example weight sets). Most score terms were recovered with mean weights greater than zero by more than one standard deviation, confirming their importance for explaining RNA structure and energetics. These terms included *stack_elec*, which models the electrostatic interaction between stacked nucleobases, an effect previously posited by several groups to be important for understanding fine-scale RNA energetics (14, 33). Terms with wider variance across weight sets could be explained through their covariance with other terms. For example, some pairs of score terms, such as the nucleobase stacking term *fa_stack* and the van der Waals term *fa_atr*, model similar physical effects, but other pairs model opposing effects in helix association, such as hydrogen bonding *hbond_sc* and the solvation term for burying polar moieties *geom_sol_fast* (SI Appendix, Table S6). The weights of these pairs varied significantly across optimized weight sets, but linear combinations of these weight pairs were nearly invariant across the weight set collection (SI Appendix, Figure S1).

Despite the variations and co-variations observed across this large collection of weight sets, each weight set gave an RMSD accuracy of better than 0.58 kcal/mol for canonical base pairs and dangling ends, with a mean accuracy of 0.40 kcal/mol across all training data. These accuracies were significantly better than RMSDs of 1.51 kcal/mol and 1.23 kcal/mol, respectively, obtained with the original structure prediction weights, supporting the need for reweighting (SI Appendix, Table S5). The RMSD over just the canonical base pairs was 0.28 kcal/mol (Figure 3a), comparable in accuracy to the initial experimental estimates of these values (10, 12) and within the estimated systematic errors of our calculation strategies (0.3 kcal/mol; see SI Appendix, Table S1 and Table S3). For the dangling-end data, RECCES-Rosetta also gave an excellent RMSD of 0.41 kcal/mol (Figure 3b). For these data, the largest deviations from experiment were tagged as having the highest expected error from weight uncertainties by RECCES, supporting this method of error computation (see, e.g., 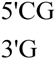 dangling end in Table 2). For both sets of NN parameters, the RMSD errors were significantly better than the range of experimental values (2.5 kcal/mol and 1.5 kcal/mol for canonical stacked pairs and dangling ends, respectively), leading to the visually clear correlations in Figure 3a-b. The terminal penalty for A-U relative to G-C was also recovered with a similar error (0.3 kcal/mol; SI Appendix includes further discussion and computation of other terminal base pair contributions).

**Figure 3.**
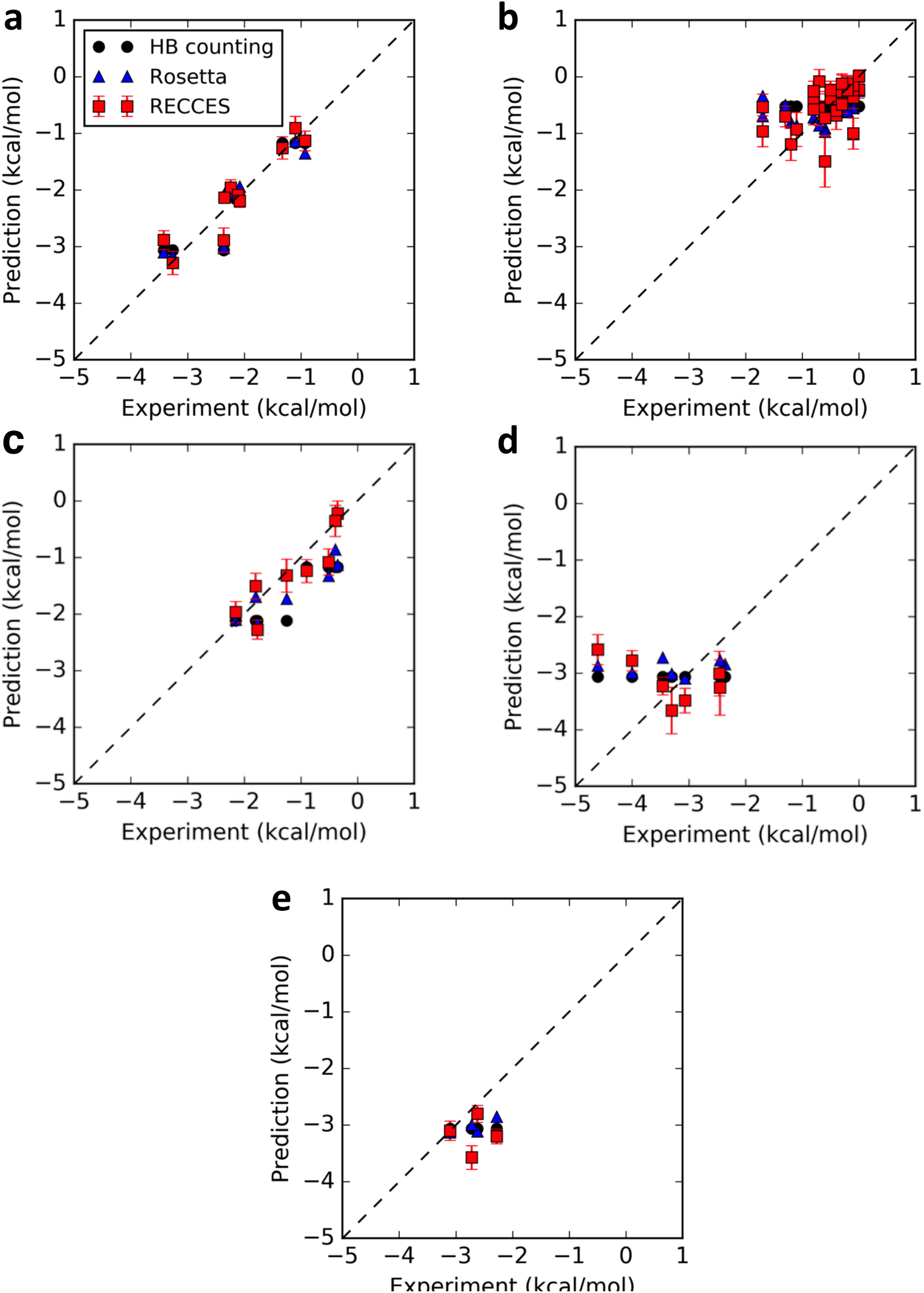
Calculations vs. experiment for each nearest neighbor parameter set. (a) Canonical stacked pairs; (b) single-nucleotide dangling ends; (c) stacked pairs including one G-U pair; (d) stacked pairs including at least one iG-iC pair; (e) stacked pairs including one D-U pair. All panels are drawn with the same axis limits and a line of equality (dashed) to aid cross-panel visual comparison.

Since we directly trained the RECCES score function against the experimental dataset, the accuracies of these results were expected. Nevertheless, we gained further confidence in the use of Rosetta-derived energy functions and RECCES framework by comparing its performance to the results of two simpler models trained on the same data. First, a three-parameter hydrogen-bond counting model, similar to simple phenomenological models that inspired the NN parametrization (34) (see SI Appendix, Supplemental Methods), achieved RMSD accuracies of 0.29 kcal/mol and 0.45 kcal/mol on canonical stacked pairs and dangling ends, respectively, slightly worse than the RECCES results (0.28 kcal/mol and 0.41 kcal/mol, respectively), despite including fitted parameters that account for conformational entropy loss of base pairs and dangling ends. Second, we evaluated a prior single-conformation Rosetta method, which uses the same energy function as RECCES-Rosetta but evaluates the score only for a minimized helix conformation (35) and fits three parameters. This model achieved accuracies of 0.30 kcal/mol and 0.44 kcal/mol for canonical stacked pairs and dangling ends, respectively, again worse than the RECCES-Rosetta results despite including separately fitted conformational entropy terms. For all three models, the largest deviation was for the stacked pair 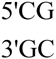, which is less stable than the other stacked pairs with two G-C pairs by 1 kcal/mol; still, even for this parameter, the RECCES-Rosetta calculations were more accurate than the simpler models. These comparisons supported the utility of the RECCES-Rosetta method compared to less generalizable models. However, the significance in the accuracy improvement was difficult to rigorously evaluate since the models contained different numbers and types of parameters; we therefore turned to independent test sets and blind predictions.

### Tests on independent nearest neighbor parameter measurements

Recent comprehensive experimental measurements have updated the NN parameters for stacked pairs involving G-U ‘wobble’ pairs next to canonical Watson-Crick pairs (11). Because these values were not used in the training of the models herein and because the geometry of G-U wobble pair is distinct from G-C and A-U pairs (Figure 1b), this set of measurements offered strong tests of modeling accuracy. Furthermore, the expected error in the RECCES-Rosetta calculations from weight uncertainties, based on variation across the large collection of weight sets, was 0.22 kcal/mol (Table 2), less than the estimated ~0.3 kcal/mol systematic error (SI Appendix, Tables S1 and S3). Both error contributions were significantly less than full range of predicted NN parameters (2.1 kcal/mol), supporting the strength of this test. The actual RMSD accuracy across these G-U NN measurements was 0.32 kcal/mol for RECCES-Rosetta, nearly as accurate as the recovery of training set stacked pairs (0.28 kcal/mol) and comparable to expected systematic errors. Furthermore, this accuracy over G-U-containing stacked pairs outperformed the RMSD values calculated from hydrogen-bond counting and single-conformation Rosetta scoring methods (0.59 and 0.49 kcal/mol, respectively) by 50-80%, supporting the importance of carrying out detailed physical simulations of the conformational ensemble via RECCES over simpler approaches. Here and below, the predictions and their estimated errors were calculated by computing means and standard deviations of NN parameters across the full collection of weight sets discovered by RECCES. Compared to this averaging over multiple models, using the single weight set with best fit to the training data gave slightly lower accuracies on the test data (SI Appendix, Table S5) (36).

A more difficult test involved seven previously measured NN parameters of a non-natural base pair, iG-iC (Figure 1d).(37) The RMSD for the iG-iC test case was 0.99 kcal/mol, mainly due to two significant outliers: 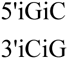 and 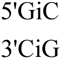 (Figure 3d). The predicted NN parameters for these outliers were larger than experimental values (less stable) by 2.2 and 1.3 kcal/mol, respectively. Nevertheless, over the other five iG-iC NN parameters, the RMSD was 0.51 kcal/mol and the discrepancies appeared primarily due to systematic offset in the predictions (Figure 3d). The accuracy was comparable to the maximum errors expected from weight uncertainties (0.4-0.5 kcal/mol) and similar, in terms of relative accuracies, to the canonical and G-U-containing stacked pairs above. Compared to RECCES-Rosetta, the simpler hydrogen-bond counting and single-conformation Rosetta scoring models gave 15-20% better accuracies (0.79 and 0.85 kcal/mol, respectively; 0.47 and 0.44 kcal/mol, excluding outliers); but both simple models gave near-constant NN parameters (range less than 0.3 kcal/mol) over all stacked pairs, providing no explanation for the 2.2 kcal/mol range in experimental measurements or for the outliers (Figure 3d). On one hand, the two outliers suggest that some important physical effect is missing or incorrectly implemented in the current calculation procedure (see Discussion below). On the other hand, the excellent accuracies over the other iC-iG-containing stacked pairs, along with the performance in the G-U test set, motivated us to continue with blind comparisons.

### Blind tests involving diaminopurine-uracil base pairs

As a blind test, we applied RECCES-Rosetta to predict the NN parameters for stacked pairs involving a distinct non-natural base pair, 2-6-diaminopurine paired with uracil (D-U, Figure 1d). Predictions of these parameters (Table 2) suggested a wide range of NN values and confirmed that errors from weight uncertainties were smaller or comparable to other systematic sources of error (0.3 kcal/mol). To test these predictions, we measured NN parameters for the four stacked pairs involving D-U next to G-C pairs, which were expected to have a range of 0.8 kcal/mol. SI Appendix, Table S7, gives construct sequences and experimental folding free energy values for these constructs; and Table 2 and SI Appendix, Table S8, summarize the NN parameter estimation. The RMSD of the RECCES-Rosetta blind predictions was 0.63 kcal/mol (Figure 3e). The hydrogen bond counting and single-conformation Rosetta scoring models, which fared worse than RECCES-Rosetta in most tests above, gave RMSDs of 0.48 and 0.40 kcal/mol, respectively, better than RECCES-Rosetta by 24-37% (Table 1). This result is similar to what we observed in the iG-iC test case; indeed, again, the two simple models produced near constant predictions (range < 0.2 kcal/mol) for the D-U stacked pairs that did not account for the 0.8 kcal/mol range of the measured values (Figure 3e). Given the blind nature of the test and our attempts to ensure its power to falsify our calculations, this test unambiguously indicated that some physical term is missing in the current Rosetta all-atom energetic model (as well as simpler models). Nevertheless, the results are encouraging: the blind predictions from each of the three models over each of four NN values separately achieved better than 1 kcal/mol accuracy compared to subsequent experimental measurements.

**Table 1.** **Accuracies of nearest neighbor parameter predictions from computational models**.

| RNA motif category | Number of motifs | RMSD accuracy (kcal/mol) |  |  |  |
| --- | --- | --- | --- | --- | --- |
|  |  | Hydrogen-bond counting | Single-conf. Rosetta | RECCES-Rosetta | RECCES-Rosetta refitted<sup>d</sup> |
| Canonical<sup>a</sup> | 10 | 0.29 | 0.30 | 0.28 | 0.41 |
| Dangling<sup>a</sup> | 32 | 0.45 | 0.44 | 0.41 | 0.43 |
| G-U<sup>b</sup> | 8 | 0.59 | 0.49 | 0.32 | 0.32 |
| iG-iC<sup>b</sup> | 7 | 0.79 | 0.85 | 0.99 | 1.08 |
| D-U<sup>c</sup> | 4 | 0.48 | 0.40 | 0.63 | 0.46 |
| All | 61 | 0.50 | 0.49 | 0.50 | 0.53 |

<sup>a</sup> Data used in training the models.<sup>b</sup> Data set aside for testing.<sup>c</sup> Blind test data.<sup>d</sup> The model was trained with all data available, so all entries in the column are training data.**Table 2. Experimental NN parameters(27, 36, 49)and RECCES-Rosetta predictions. All values in kcal/mol.**

### Post hoc fit across all data

While post hoc tests of models on prior collected data are less rigorous than blind trials, they can help guide future work. As a final test of our study, we wished to understand possible explanations for the worse accuracy of RECCES-Rosetta in iG-iC test cases and blind D-U trials compared to the G-U test cases. One model for this inaccuracy was that overfitting of energy function weights to the training data worsened predictive power over the new data. Another (not necessarily exclusive) model was that the underlying energy function derived from Rosetta score terms was fundamentally incapable of modeling the available NN data under any weight set with the RECCES procedure. We were able to test these models by carrying out a post hoc global fit of energy function weights over all available NN data ((Figure 4 and Table 1). As expected, we observed better fits to the test data, including an improvement in RMSD accuracy for the four D-U stacked pairs from 0.63 kcal/mol to 0.46 kcal/mol; this suggests a modest overfitting to the training set in the studies above. However, we observed somewhat worse fits to the training data, including a worsening of RMSD accuracy for the ten canonical stacked pairs from 0.28 to 0.41 kcal/mol, worse than expected systematic errors in our calculations (0.3 kcal/mol; SI Appendix, Tables S1 and S3) supporting the second model of fundamental energy function inaccuracy. Furthermore, this global fit still failed to account for the two striking > 1 kcal/mol outliers involving iG-iC base pairs, giving evidence for the second model: energetic calculations based on the Rosetta score function are fundamentally incapable of accounting for all the data within expected error, even with a post hoc optimized weight set.

**Figure 4.**
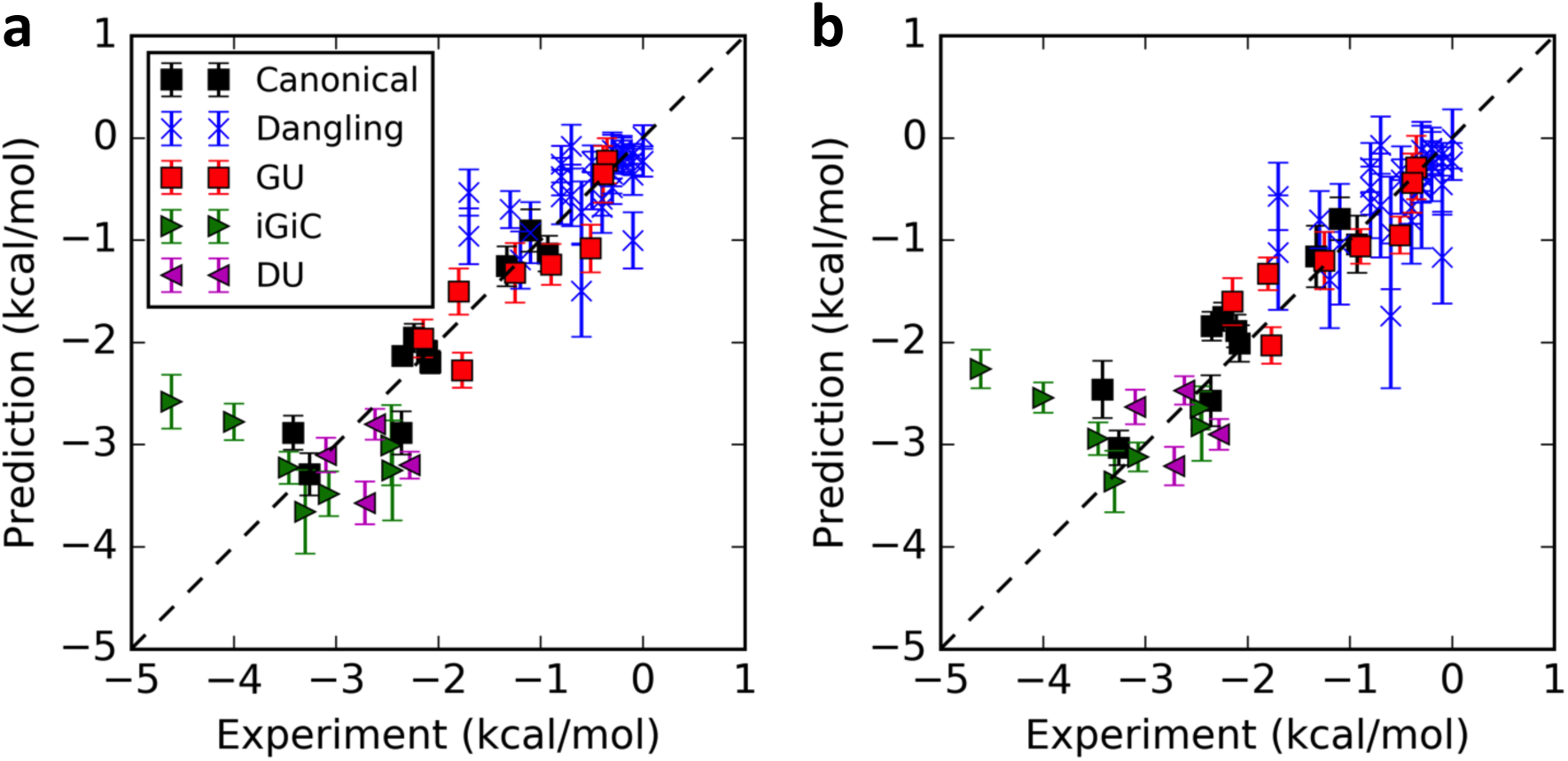
Calculations vs. experiment across all nearest neighbor parameters. Comparisons are based on (a) RECCES-Rosetta weight sets trained on canonical and dangling end data (same values as Figure 3) and (b) ‘best-case’ weight sets fitted *post hoc* over all available NN parameters, including D-U stacked pairs measured for blind predictions.

## Discussion

This study reports the first blind test of the predictive power of high-resolution, all-atom modeling methods for RNA folding energetics. We developed a Reweighting of Energy-function Collection with Conformational Ensemble Sampling (RECCES) strategy in the Rosetta framework that rigorously models conformational ensembles of single strand and helical states, is computationally efficient (hours with currently available CPUs), and brackets systematic errors based on comprehensive reweighting tests. When compared to simpler phenomenological methods, RECCES-Rosetta achieved excellent RMSD accuracies for the nearest neighbor parameters of canonical base pairs, dangling ends, and G-U pairs, but somewhat worse results for NN parameters involving non-natural base pairs iG-iC and D-U. The latter D-U parameters were measured after the predictions as a blind test. The computational accuracies were better than 1 kcal/mol in all cases, based on RMSD values over each separate set of NN parameters (0.28, 0.41, 0.32, 0.99, and 0.63 kcal/mol for canonical, dangling end, G-U, iG-iC, and D-U parameters, respectively) and also individually for each of the four blind predictions. This accuracy level is significantly smaller than the 2-3 kcal/mol ranges measured for these sets of NN values ( Figure 4, Table 2), is comparable to errors in *ad hoc* fits used in the current nearest-neighbor model for most motifs (25–28), and is generally better than molecular dynamics calculations that remain significantly more expensive (21, 22). The generality of the RECCES-Rosetta framework and this level of success in initial tests supports the further development of RECCES-Rosetta for non-natural nucleotides and for motifs more complex than the helical stacked pairs and dangling ends considered herein.

While achieving consistently sub-kcal/mol accuracies, there is room for improvement in the RECCES-Rosetta approach. For example, the modeling does not account for the 1 kcal/mol stability increase of the 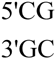NN parameter relative to 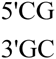; the electrostatic term *stack_elec* does favor the former, but is not assigned a strong enough weight in the final fits to account for the stability difference. Also, the RMSD accuracies still remain larger than estimated systematic errors (0.3 kcal/mol), particularly for the non-natural base pairs in the test data, and the discrepancies remain even if those data are included in a *post hoc* fit of the energy function weights to all available measurements. Our results help bracket which strategies might improve the accuracy and which might not. On one hand, non-natural pairs present their atomic moieties in different bonded contexts, which might modulate the strengths of hydrogen bonds or other interactions that they form. For example, a previous analysis suggested that the hydrogen bonds in an iG-iC base pair might be stronger than in a G-C base pair by approximately 0.4 kcal/mol (38). Accounting for this effect would be predicted to offset our calculated NN parameters for all iG-iC stacked pairs, without changing their relative ordering, and cannot account for strong ‘outliers’. Indeed, if we added an extra fitting term for stabilizing iG-iC pairing, the RMSD accuracy over these data did not significantly improve (0.96 kcal/mol, vs. 0.99 kcal/mol without the extra term). On the other hand, several unmodeled factors are sensitive to the ordering of base pairs within stacked pairs and could affect the relative ordering of NN parameters within each set. For example, the current Rosetta all-atom score function models electrostatics through fixed charges with a distance-dependent dielectric and does not explicitly model water or counterions that may differentially stabilize the base pair steps [(39); see, however, (40)]. Recent and planned additions of nonlinear Poisson-Boltzmann solvation models, polarizable electrostatic models, and a potential of mean force for water-mediated hydrogen bonding into the Rosetta framework should allow evaluation of whether these physical effects can improve accuracy of nearest neighbor parameter calculations to the 0.3 kcal/mol fundamental limit of the RECCES method. If these models can also be expanded to calculate the temperature dependence of solvation, it may also become possible to compare calculated and measured entropies and enthalpies of the NN parameters, which are well-measured but may be dominated by solvation effects. In addition, we propose that calculations for recently characterized stacked pairs that give anomalous NN parameters, including some tandem G-U stacked pairs (11) and pseudouridine-A-containing stacked pairs (41, 42), could offer particularly stringent tests.

Continuing work in modeling RNA energetics will benefit from further blind trials, perhaps in a community-wide setting analogous to the ongoing RNA-puzzles structure prediction trials (43, 44). The prediction of two kinds of parameters could serve as future blind tests. First, based on the results herein, non-natural base pairs offer good test cases and require the same amount of computational power as canonical base pair NN parameter estimation. Alternative approaches based on, e.g., molecular dynamics, should also be applicable to these cases. We have completed RECCES-Rosetta predictions for additional stacked pairs involving iG-iC and D-U pairs, as well as for inosine-cytosine (I-C) base pairs (Figure 1e; Table 2), but are waiting to make experimental measurements until there are comparison values from other groups and approaches. Second, future blind trials might involve predicting energetics of RNA motifs more complex than those considered herein, such as apical loops, internal loops, multi-helix junctions, and tertiary interactions. For these cases, an expansion of the RECCES approach in which physically realistic candidate conformations of each motif are first estimated with structure prediction (35, 45) and then subjected to rigorous RECCES-based free energy calculations, may offer predictive power. Such an approach may also allow calculations of next-nearest-neighbor effects and development of rapid approximations to estimate conformational entropy of candidate conformations, which would be useful for structure prediction and design (see also SI Appendix, Figure S2). A new generation of high-throughput RNA biochemistry platforms (46–48) offers the prospect of both training these next-generation energetic prediction algorithms and carrying out blind tests with many thousands of measurements.

## Materials and Methods

Details of nearest-neighbor parameter estimation with RECCES (including basic equations, simulation parameters, and energy function) and with simple single-conformation methods, as well as methods used to experimentally estimate nearest neighbor parameters for helices with D-U base pairs, are presented in the SI Appendix.

## Acknowledgements

We thank J. Yesselman for help in generating the partial charges for non-natural RNA bases and P. Sripakdeevong, K. Beauchamp, W. Greenleaf, and members of the Das lab for useful discussions. This work is supported by an HHMI International Student Research Fellowship (FC), a Stanford BioX graduate student fellowship (FC), a Burroughs-Wellcome Career Award at Scientific Interface (RD), and NIH grant NIGMS R21 GM102716 (RD). The calculations were performed using the TACC Stampede cluster through an XSEDE allocation (project MCB120152), and the Stanford BioX^3^ cluster.

